# Central metabolism is a key player in *E. coli* biofilm stimulation by sub-MIC antibiotics

**DOI:** 10.1101/2022.09.14.507886

**Authors:** Luke N. Yaeger, Shawn French, Eric D. Brown, Jean Philippe Côté, Lori L. Burrows

## Abstract

Exposure of *Escherichia coli* to sub-inhibitory antibiotics stimulates biofilm formation through poorly characterized mechanisms. Using a high-throughput Congo Red binding assay to report on biofilm matrix production, we screened ∼4000 *E. coli* K12 deletion mutants for deficiencies in this biofilm stimulation response. Mutants lacking *acnA, nuoE*, or *lpdA* failed to respond to sub-MIC novobiocin, implicating central metabolism and aerobic respiration in biofilm stimulation. These genes are members of the ArcA/B regulon – controlled by a respiration-sensitive two-component system. Mutants of *arcA* and *arcB* had a ‘pre-activated’ phenotype, where biofilm formation was already high relative to wild type in vehicle control conditions and failed to increase further with the addition of sub-MIC antibiotics. Supporting a role for respiratory stress, the biofilm stimulation response was inhibited when nitrate was provided as an alternative electron acceptor. Deletion of genes encoding the nitrate respiratory machinery abolished its effects, and nitrate respiration increased during growth with sub-MIC antibiotics. In probing the generalizability of biofilm stimulation, we found that the stimulation response to translation inhibitors was minimally affected by nitrate supplementation. Finally, using a metabolism-sensitive dye, we showed spatial co-localization of increased respiration with sub-MIC bactericidal antibiotics. By characterizing the biofilm stimulation response to sub-MIC antibiotics at a systems level, we identified multiple avenues for design of therapeutics that impair bacterial stress management.

## Introduction

Biofilms are adherent bacterial communities and a common form of bacterial growth (Flemming & Wuertz, 2019; Serra & Hengge, 2021). They also play a significant role in infection and antibiotic tolerance (Brooun, Liu, & Lewis, 2000; Costerton et al., 1987; Donlan & Costerton, 2002). Biofilms consist of bacterial cells surrounded by a self-produced matrix of exopolysaccharides, extracellular DNA (eDNA), and proteinaceous adhesins that provide protection against physical and chemical threats (Nadell, Xavier, & Foster, 2009). The composition of the *Escherichia coli* biofilm matrix varies among strains, but generally includes a combination of curli fimbriae (an amyloid), eDNA, and the exopolysaccharides colanic acid, cellulose, and poly-N-acetyl glucosamine (PNAG) (Agladze, Wang, & Romeo, 2005; Hung et al., 2013; Claire Prigent-Combaret et al., 2000; Wu & Xi, 2009). Curli, PNAG, the autotransporter antigen 43, and type 1 fimbriae all contribute to structural integrity of *E. coli* biofilms through cell-cell and cell-surface adhesion (Danese, Pratt, Dove, & Kolter, 2000; Harris et al., 1990; Claire Prigent-Combaret et al., 2000; Wang, Preston, & Romeo, 2004). While specific pro-biofilm components drive the emergent properties of biofilms, the production of these components is influenced by an array of central pathways in *E. coli* (Smith et al., 2017). The *E. coli* biofilm life cycle is well-studied, involving reversible and irreversible attachment, maturation, and dispersal (C. Beloin, Roux, & Ghigo, 2008). Although biofilms form naturally, growth conditions can negatively or positively influence biofilm formation (Karatan & Watnick, 2009).

Treatment with sub-<u>m</u>inimal inhibitory <u>c</u>oncentrations (sub-MIC) of antibiotics stimulates biofilm formation (M. R. Ranieri, Whitchurch, & Burrows, 2018). Biofilms are stimulated by multiple classes of antibiotics, in many different organisms (Kaplan, 2011), suggesting there are conserved responses to antibiotic stress that converge on increased biofilm formation. The diverse stressors that trigger biofilm formation extend beyond antibiotics to type six secretion system effectors, bacteriocins, bacteriophages, and biocides (Gödeke, Paul, Lassak, & Thormann, 2011; Lories et al., 2020; Oliveira et al., 2015; Shemesh, Kolter, & Losick, 2010a). This list of triggers informed the idea of competition sensing, where cells are proposed to sense damage or kin cell death and respond to protect themselves (Cornforth & Foster, 2013; Le Roux et al., 2015; Ting et al., 2022). The induction of biofilms by sub-MIC antibiotics and the ability of biofilms to tolerate multiple stressors suggests that this phenotype falls under the umbrella of competition sensing. Despite the importance of antibiotics, interspecies interactions, and biofilm formation in *E. coli* ecology, we know relatively little about how antibiotic stress generates a physiological signature that is relayed into increased biofilm formation.

We view biofilm stimulation as a defensive response to stressors (M. R. Ranieri et al., 2018) and consistent with this hypothesis, major *E. coli* stress response pathways Cpx, Rcs, and OmpR influence *E. coli* biofilm formation (Ferrières & Clarke, 2003; Otto & Silhavy, 2002; C. Prigent-Combaret et al., 2001). There is also significant input of the stress-responsive sigma factor RpoS and oxidative stress responses in biofilm regulation (Gambino & Cappitelli, 2016; Mika & Hengge, 2014; Smith et al., 2017). To better understand the mechanism underlying the response to sub-MIC antibiotics, we applied a systems genetics approach to identify genes involved in biofilm stimulation. Here we describe a 1536 colony density, high-throughput screen of the Keio collection – a single gene knockout library of *E. coli* K12 (Baba et al., 2006) – that used uptake of the curli, cellulose, and PNAG-binding dye Congo Red as a readout to identify genes important for biofilm stimulation in response to sub-MIC antibiotics of three different classes: cefixime (CEF), novobiocin (NOVO), and tetracycline (TET). This screen identified multiple genes in central metabolism and respiration as important for biofilm stimulation. Follow-up experiments also implicated the two-component system ArcA/B in this response, suggesting that oxidative stress was a key trigger. Supporting this idea, provision of nitrate in aerobic growth conditions inhibited biofilm stimulation by certain sub-MIC antibiotics. We linked this observation to nitrate’s role as an alternative electron acceptor using nitrate respiratory mutants and a nitrate reduction assay. Finally, we showed that increased respiratory activity spatially colocalizes with sub-MIC antibiotics. These results suggest that perturbations of metabolism and respiratory chain activity resulting from exposure to sub-MIC antibiotics drives biofilm stimulation.

## Results

### Development of a high-throughput 1536-colony density biofilm stimulation screen

The goal of this study was to identify genes and pathways involved in the *E. coli* biofilm stimulation response using a systems approach. First, we used a 96-well peg-lid assay to measure *E. coli* biofilm formation and identified antibiotics that elicited a robust stimulation response. As expected, biofilm formation spiked at concentrations below those that decreased planktonic growth (**Figure 1a**). From the panel of antibiotics tested (**Supplementary Figure S1**), we selected CEF, NOVO, and TET as three model antibiotics with distinct mechanisms of action (targeting peptidoglycan synthesis, DNA replication, and protein synthesis, respectively) that stimulate robust biofilm formation. Although the peg-lid assay is a reliable method for quantifying biofilms, it is low throughput and less amenable to large-scale genetic screens.

**Figure 1.**
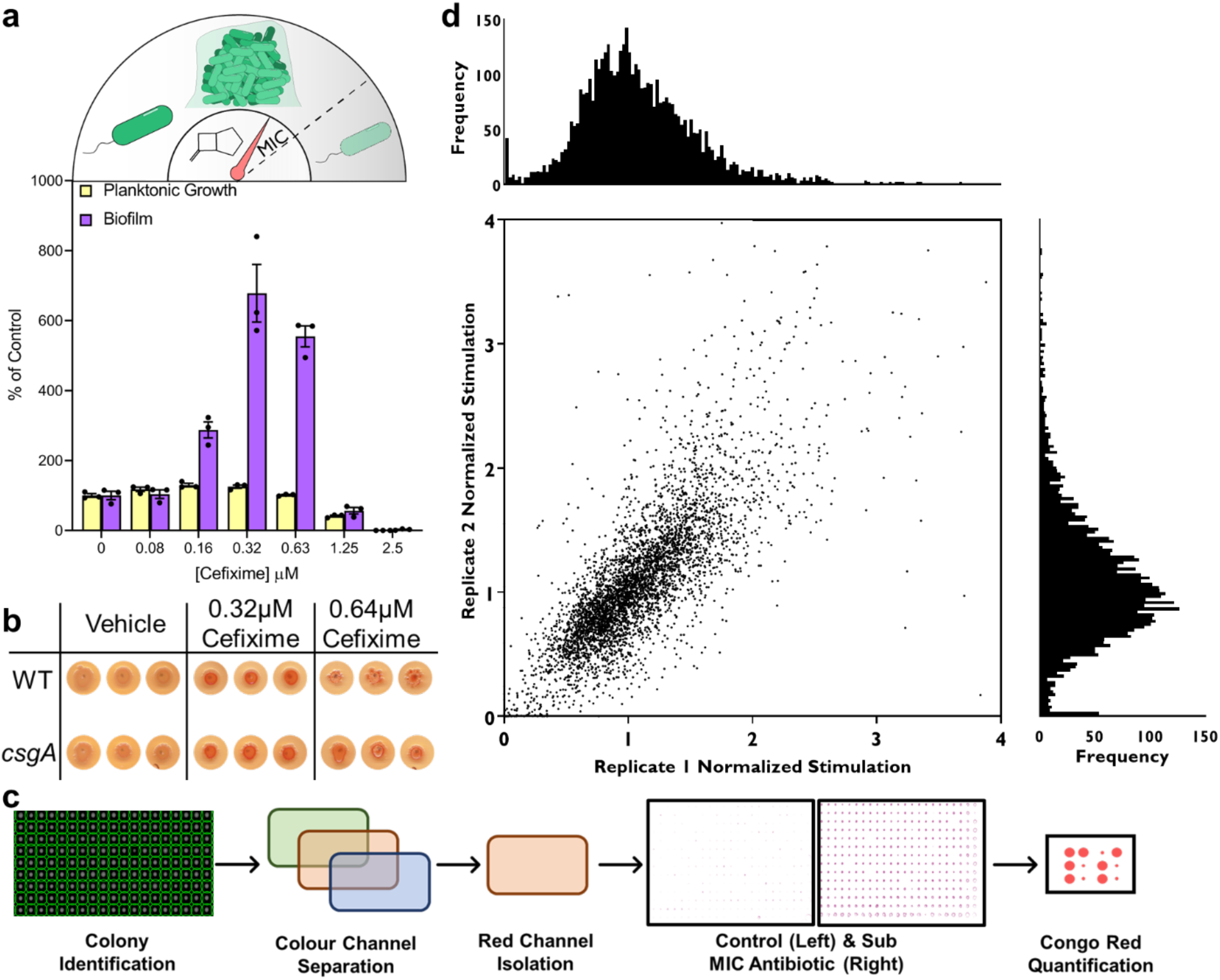
A high throughput screen for genes involved in biofilm stimulation. **a)** (top) An illustration of the biofilm stimulation response, where tuning the levels of antibiotics to near the MIC stimulates biofilm formation. (bottom) A 96-well peg lid assay to quantify biofilm stimulation in a dose response for *E. coli* K12 BW25113. Values are shown as a percent of the vehicle control and each point of a technical triplicate are shown with the bars showing the mean and standard error of the mean shown with lines. Yellow bars show planktonic growth and purple bars show biofilm. The data are representative of at least 3 biological replicates. **b)** Measuring biofilm formation using Congo Red on solid 50:50 agar plates with and without sub-MIC CEF. Colonies are shown for the WT K12 and the *csgA* mutant from the Keio collection **c)** Screening workflow for finding mutants deficient in biofilm stimulation. **d)** Screening results for CEF shown in a replica plot with extremely high stimulation data points (above 4) removed to help visualize the low stimulation hits. Histograms show the distribution of values across each replicate shown on their respective axes.

To create a colony-based biofilm assay for high-throughput screening, we leveraged the ability of the dye Congo Red (CR) to bind the *E. coli* extracellular matrix components curli, cellulose, and PNAG (Carzaniga, Antoniani, Dehò, Briani, & Landini, 2012; Serra, Richter, & Hengge, 2013). We spotted *E. coli* K12 BW25113 on 50:50 LB:PBS agar + CR supplemented with sub-MIC (1/4 to 1/2 MIC) CEF, NOVO, or TET, and grew them for 24 h at 37ºC – generally considered a suppressive media and temperature for curli expression and CR binding (Gualdi et al., 2008; Olsen, Arnqvist, Hammar, & Normark, 1993). While the resulting colonies on antibiotic-free control plates showed no colour change, those on sub-MIC antibiotic-supplemented plates bound CR and turned red (**Figure 1b**). This result indicates that treatment with sub-MIC antibiotics allows *E. coli* to overcome the suppressive effects of the growth conditions on CR binding. Interestingly, curli are likely not responsible for antibiotic-induced CR binding, as a *csgA* mutant still showed increased staining. Cellulose and PNAG are the other CR-binding *E. coli* biofilm components; however, K12 strains are unable to produce cellulose (Serra et al., 2013). Therefore, the only known CR binding polymer remaining in the *csgA* mutant is PNAG, an exopolysaccharide reported previously to be important for translation inhibitor-induced biofilm formation (Boehm et al., 2009).

We then endeavoured to identify the physiological changes that precede production of biofilm components. The increase in CR binding served as a proxy for biofilm stimulation in our high throughput screen, as highlighted in **Figure 1c**. Briefly, the Keio collection was arrayed in 1536 density on CR-supplemented agar plates, without and with sub-MIC NOVO, and grown for 24 h at 37ºC, then imaged and analyzed with Fiji (Schindelin et al., 2012). Growth was quantified by measuring colony density; then, the colour channels for each image were separated to isolate the red channel, and the red pixel intensity measured to quantify CR binding. To correct for plate positioning effects, CR binding and growth were normalized by the interquartile mean for each colony’s respective column and row (Mangat, Bharat, Gehrke, & Brown, 2014). We generated a CR enrichment value by dividing the normalized CR binding value by the normalized growth value, and plotted the enrichment values (i.e. normalized stimulation) for both replicates (**Figure 1d**). Mutants with enrichment values at least two standard deviations below the mean in both replicates were classified as hits. The screen also identified mutants with enrichment values well above average. These high values could indicate hyper-biofilm stimulation mutants or be the result of mutations causing small colony sizes, high basal CR binding, or slight changes in antibiotic sensitivity that decreased colony size or impacted the relative MIC, to result in more biofilm without affecting a central stress response pathway. For these reasons, we did not investigate the high enrichment-value mutants as they were more liable to be false positives.

### Central metabolism and respiration genes as major components of biofilm stimulation

We verified loss of biofilm stimulation in response to sub-MIC antibiotics for the hits from the screen using a dose-response peg lid assay (**Figure 2b and c**). The maximal level of biofilm stimulation for deletion mutants of *acnA* (encoding aconitase), *lpdA* (encoding lipoamide dehydrogenase), *nuoE* (encoding subunit of complex I in the electron transport chain), *lpp* (encoding Braun’s lipoprotein), and *ydcU* (encoding a predicted polyamine transporter) was markedly reduced upon treatment with sub-MIC NOVO or CEF compared to the wild type.

**Figure 2.**
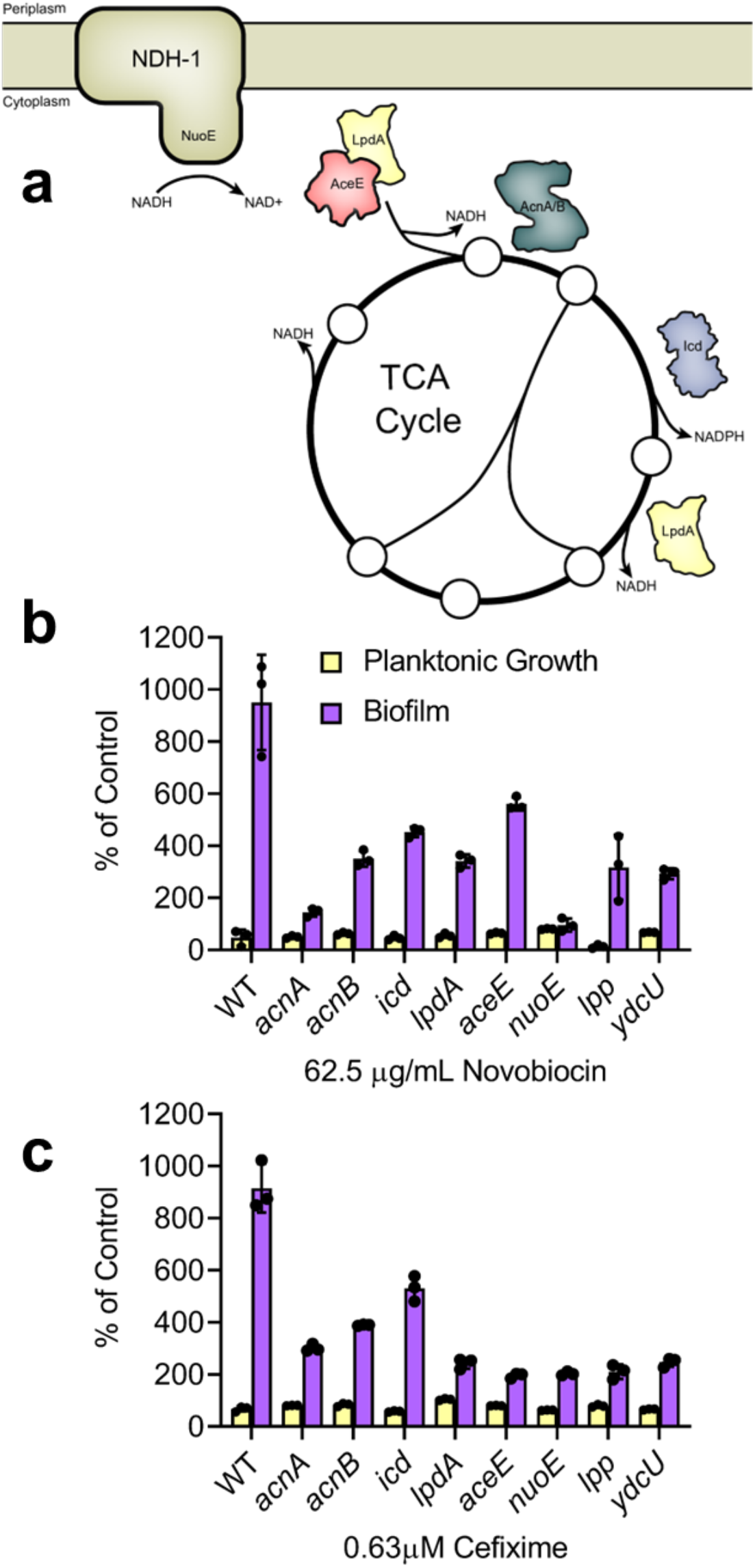
Verification of novobiocin screening hits. **a)** Hits lacking genes involved in central metabolism are illustrated to show their location in the TCA cycle and respiration. Circles indicated metabolites of the TCA cycle. Growth and biofilm formation were measured for cells grown in 50:50 LB with serial dilutions of **b)** NOVO or **c)** CEF using peg-lid assays. Circles on the graphs show data points of technical triplicates that are representative of two biological replicates. Data are shown as a percent of control relative to each mutant’s respective untreated control. Bars indicate the technical triplicate mean and the error bars show the standard error of the mean.

The hits were enriched for mutants in central metabolism and respiration (**Figure 2a**), consistent with previous studies suggesting that antibiotics alter metabolic state and respiration as a part of their lethality (Belenky et al., 2015; Dwyer et al., 2014; Lobritz et al., 2022, 2015; Stokes, Lopatkin, Lobritz, & Collins, 2019). Thus, we focused on *acnA, acnB, lpdA*, and *nuoE* for further examination because of their common roles in central carbon metabolism and aerobic respiration (Karp et al., 2018). Notably, Mathieu *et al*. found that *acnB* was upregulated in response to many different sub-MIC antibiotics (Mathieu et al., 2016), while *nuo* mutants were implicated in *E. coli* stress-induced mutagenesis via ubiquinone pool sensing by the ArcA/B two-component system (Al Mamun et al., 2012). To ensure that we identified genes involved in central pathways for biofilm stimulation, we next re-screened the entire library with sub-MIC NOVO, CEF, and TET (CEF replica plot shown in **Figure 1**, NOVO and TET replica plots shown in **Supplementary Figure S2**). We selected hits from those screens with normalized enrichment values below 0.5, in both replicates, for all three antibiotics. Those with normalized growth that fell below 0.2 were excluded due to the potential for small colonies to generate false positives. All hits with their predicted functions are listed in **Supplementary Table S2**.

Using 3 different antibiotics, we identified common stimulation-deficient mutants in the lower TCA cycle, (*lipB, sucA/B, sdhB/D*) that convert 2-oxoglutarate to fumarate while generating NADH for aerobic respiration. A set of genes linked to iron-sulfur cluster biogenesis (*tusA, fdx, cysJ, cysA*) was also identified. Other hits with links to respiration included *nuoM, menC, pntB*, and *pyrD*. Mutants in ubiquinone synthesis were enriched among the hits; however, they had severe growth defects and were filtered from the final hit list. **Figure 3a** shows a STRING network interaction map for the related hits from the three-drug screen (Szklarczyk et al., 2021). Together, these data suggest that interrupting flux through the TCA cycle and respiratory chain prevents biofilm stimulation by sub-MIC antibiotic stress. Antibiotic stress was previously shown to perturb central metabolism and aerobic respiration by increasing flux through those pathways (Belenky et al., 2015; Dwyer et al., 2014; Lobritz et al., 2022, 2015; Stokes et al., 2019). In certain conditions, increased flux through the respiratory chain can generate reactive oxygen species (ROS) (Seaver & Imlay, 2004), as electrons stall on solvent-exposed flavins and are transferred to oxygen. Intracellular superoxide and hydrogen peroxide then oxidize iron-sulfur clusters, increasing the free iron available to participate in the Fenton reaction (Jang & Imlay, 2007; Keyer & Imlay, 1996). The role of iron in respiration and ROS generation is in line with the data from our screen, connecting the metabolism and iron-sulfur cluster networks shown in **Figure 3a**.

**Figure 3.**
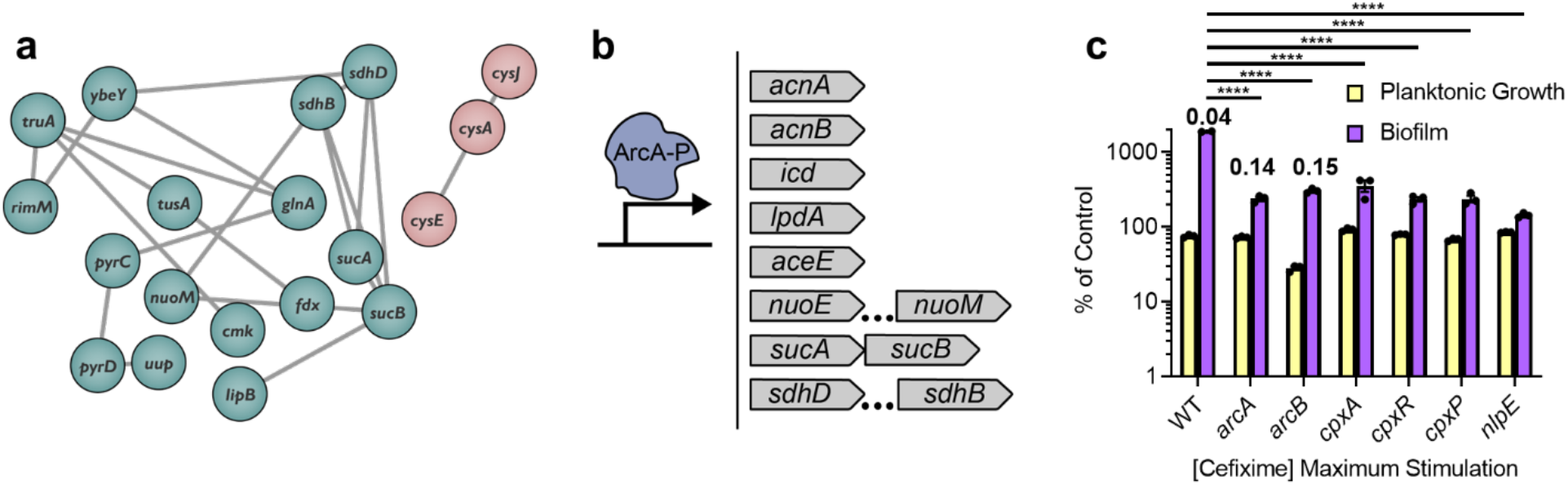
Involvement of metabolic genes and regulatory systems in biofilm stimulation. **a)** A STRING network map of connected hits from the three-drug screen. Connections between each node indicate an experimentally-determined interaction, gene fusion, neighbourhood, or co-occurrence, protein homology, co-expression, or text mining. The hits clustered into two groups, coloured green and pink. The network map was visualized in Cytoscape (Shannon et al., 2003). **b)** A cartoon depicting the binding of phosphorylated ArcA to a promoter region. The genes from our hit list regulated by ArcA are listed on the right. Ellipses indicate genes in an ArcA-regluated operon that are not directly adjacent. **c)** Biofilm peg-lid assays were done with increasing concentrations of CEF. Planktonic growth and biofilm formation are shown as a percent of the vehicle control. Bars indicate the triplicate mean and error bars indicate the standard error of the mean. Average background subtracted crystal violet absorbance values for WT, *arcA*, and *arcB* untreated condition are shown above each respective biofilm bar. The y-axis is shown as a log scale. A one-way ANOVA followed by a Dunnett’s multiple comparisons test was used to calculate statistical significance between the WT and mutant biofilm; ^****^ = p value<0.0001

We next considered how *E. coli* might sense changes in respiration state (Price, Román-Rodríguez, & Boyd, 2021). An obvious candidate was ArcA/B, a two-component regulatory system that responds to changes in respiration based on the redox state of the quinone pool, and coordinates expression of metabolism and respiration genes, including many of our hits (**Figure 3b**) (Alvarez, Rodriguez, & Georgellis, 2013). To test the involvement of ArcA/B in the response to sub-MIC antibiotics, we measured biofilm stimulation for *arcA* and *arcB* mutants (**Figure 3c**). Both had increased baseline biofilm formation relative to the parent strain (raw crystal violet absorbance values are labelled) and only minimal increases in biofilm levels in response to sub-MIC CEF exposure. These data suggest that inactivation of ArcA/B phenocopies the biofilm stimulation response. It is worth noting that a minimal level of biofilm stimulation was still possible in these mutants, suggesting that additional factors contribute to the stimulation response independent of ArcA/B.

While the ArcA/B system controls expression of many genes involved in metabolism and respiration and loss of ArcA reduces competitiveness in a biofilm (Junker, Peters, & Hay, 2006), it does not directly control genes that encode for biofilm components. Crosstalk between the Arc and Cpx systems under sub-MIC gentamicin stress was previously reported (Ćudić, Surmann, Panasia, Hammer, & Hunke, 2017), so we tested mutants in various steps of the Cpx stress response pathway for biofilm stimulation. The Cpx pathway is responsive to cell envelope stress, affects regulation of respiration complexes, and is implicated in biofilm formation (Guest, Wang, Wong, & Raivio, 2017; Otto & Silhavy, 2002). We tested mutants of *cpxA*/*P*/*R* and *nlpE* of the Cpx system. All three *cpx* mutants exhibited reduced biofilm stimulation, while *nlpE* showed no stimulation at all (**Figure 3c**). These mutants were not identified in our original screen, perhaps due to differences between the CR and peg-lid assays, or changes in antibiotic sensitivity in mutants lacking this system. CpxR/A and NlpE were implicated in *E. coli* surface sensing and biofilm formation (Christophe Beloin et al., 2004; Ma & Wood, 2009; Otto & Silhavy, 2002) and the Raivio group proposed a model in which the Cpx system detects improper assembly of complex I of the electron transport chain (ETC) while also controlling its expression (Guest et al., 2017; Tsviklist, Guest, & Raivio, 2022). Disrupting the Cpx response could suppress biofilm stimulation by dysregulating maintenance of the ETC.

### Controlling biofilm stimulation with the terminal electron acceptor nitrate

The redox state of ubiquinone/ubiquinol and the ratio of ubiquinone to menaquinone control ArcA/B activity (Alvarez et al., 2013). We reasoned that if sub-MIC antibiotics were inducing aerobic respiration changes that influenced ArcA/B activity, then supplementing an additional terminal electron acceptor could change the abundance or redox state of quinones to supress biofilm stimulation. Through these quinones, the ArcA/B system is responsive to the availability of nitrate as an electron acceptor under restricted oxygen conditions (Iuchi, Aristarkhov, Dong, Taylor, & Lin, 1994). As predicted, potassium nitrate suppressed biofilm stimulation in a dose-dependent manner without affecting the CEF MIC (**Figure 4a**). To rule out the effect of potassium cations, we tested sodium nitrate, which also inhibited biofilm stimulation (**Supplementary Figure S3)**. *E. coli* dissimilative nitrate respiration follows the ammonification pathway, where nitrate is reduced to nitrite, then ammonia, although the latter step is less frequent due to repression of nitrite reductase by nitrate and efflux of nitrite (Cole, 1996). Since aerobic conditions generally repress nitrate reduction, we controlled for nitrate’s potential ability to inhibit biofilm stimulation through pathways other than respiration. We tested whether a mutant lacking NarG, a subunit of the major nitrate reductase (Bonnefoy & Demoss, 1994), responded to nitrate supplementation. Consistent with our hypothesis that nitrate inhibits biofilm stimulation through its role as an electron acceptor, sub-MIC CEF stimulated biofilm formation by the *narG* mutant, regardless of nitrate addition (**Figure 4b**). We also directly tested if sub-MIC cefixime stimulates nitrate reduction using an orthogonal, nitrite-driven diazotization reaction that produces an azo dye (**Figure 4c**) and found a significant spike in nitrate reduction at ½ MIC (**Figure 4d**) (Buxton, 2011; Griess, 1858). These results are in line with reports of upregulation of anaerobic respiration genes by sub-MIC antibiotics in *Listeria monocytogenes* (Knudsen, Fromberg, Ng, & Gram, 2016).

**Figure 4.**
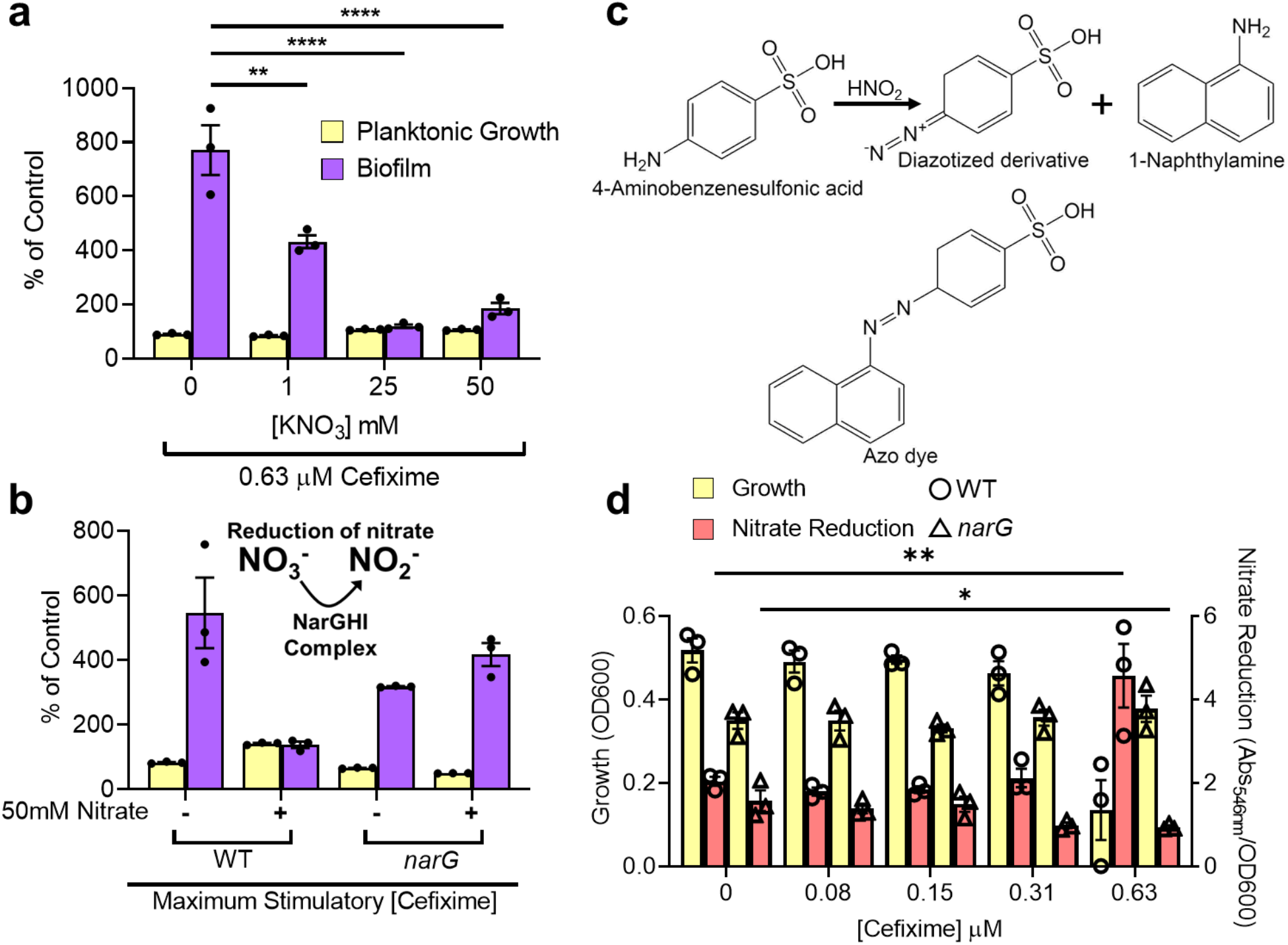
Nitrate respiration suppresses biofilm stimulation. **a)** A dose response assay showing the effect of nitrate on biofilm stimulation. The data were taken from CEF dose-response peg-lid assays and the maximum biofilm from each nitrate concentration is shown. One representative experiment from three similar biological replicates is shown. A two-way ANOVA was performed in GraphPad Prism to compare biofilm formation to the no nitrate control (** = p value <0.01, and **** = p value <0.0001) **b)** Effects of nitrate on a nitrate reductase mutant. An illustration of nitrate to nitrite reduction is shown above. Plus and minus signs indicate the presence of 50 mM nitrate in the 50:50 medium. The maximum biofilm stimulation from a dose response assay for each strain and condition is shown. Percent of control is relative to the untreated control for each strain and condition. The graph is representative of two biological replicates. **c)** The reaction scheme for detecting nitrite from nitrate reduction using a diazotization assay. Nitrite is converted to nitrous acid with acetic acid, which reacts with 4-aminobenzenesulfonic acid to create a diazonium salt that reacts with 1-napthylamine to produce a dye. Chemical structures were created in ChemDraw. **d)** Using the diazotization assay from c) to measure nitrate reduction after growth of WT (circles) or *narG* (triangles) in increasing sub-MIC CEF concentrations. Planktonic growth is plotted on the left y-axis as the sterility control subtracted OD_600_ values, and nitrate reduction is shown as the absorbance at 546 nm with sterility control subtracted, divided by the growth of the corresponding well, and plotted on the right y-axis. The graph is representative of two biological replicates. A one-way ANOVA followed by Dunnett’s multiple comparisons test was used to compare nitrate reduction between the treated wells and untreated control for each strain. There was no significant change for any condition except the nitrate reduction for the 0.63μM CEF treated WT and *narG* (* = p value <0.05, ** = p value <0.01). For all graphs, bars indicate the triplicate mean and error bars show the standard error of the mean.

### Nitrate insensitivity reveals divergence of translation inhibitors in biofilm stimulation pathway

Interestingly, we found that adding nitrate suppressed biofilm stimulation by NOVO but not TET (**Supplementary Figure S4**). To probe the basis of TET insensitivity to nitrate, we tested mulitple translation inhibitors with different features, to rule out TET-specific effects (**Figure 5**). TET acts on the 30S subunit of the ribosome by blocking binding of charged tRNAs to the A site and inhibits translation initiation (Nakahigashi et al., 2016; Wilson, 2009).

**Figure 5.**
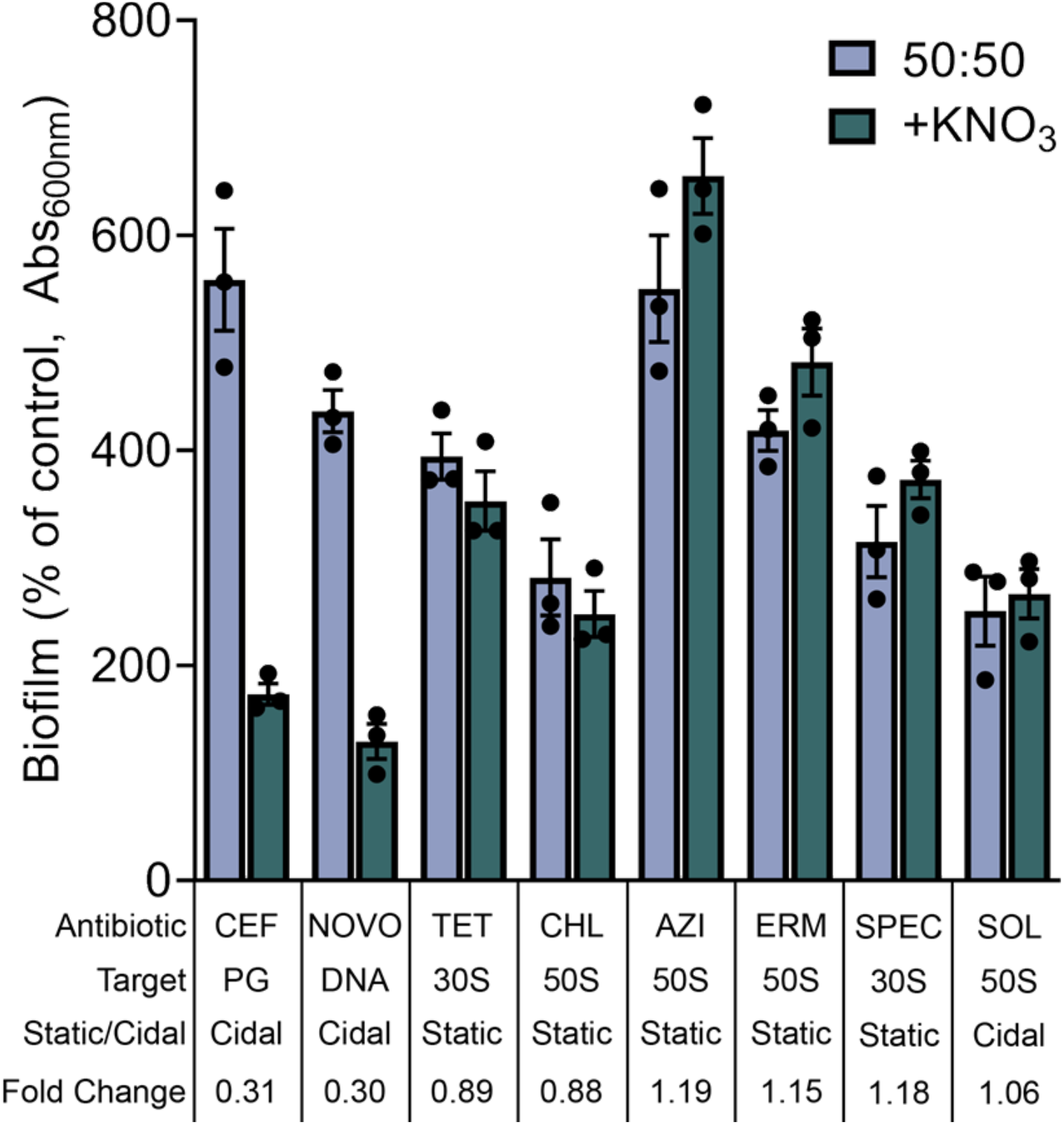
Differing effects of nitrate on biofilm stimulation across a panel of antibiotics. Biofilm assays were performed using peg lids in 50:50 media with or without 50mM potassium nitrate. Each antibiotic (CEF=cefixime, NOVO=novobiocin, TET=tetracycline, CHL=chloramphenicol, AZI=azithromycin, ERM=erythromycin, SPEC=spectinomycin, SOL=solithromycin) was tested across a range of 2x serial dilutions as before, and the maximum biofilm stimulation (highest average at one concentration) for each condition was plotted as a percent of the untreated control. “30S” and “50S” indicate the target subunit for ribosome inhibiting compounds. “Static” and “Cidal” are shorthand for bacteriostatic and bactericidal, respectively. Fold change was calculated by dividing the maximum biofilm stimulation value of the nitrate treated sample by the no nitrate control value. The data are representative of three biological replicates.

Chloramphenicol (CHL) targets the 50S subunit at the peptidyl transferase center and inhibits peptide bond formation (Wilson, 2009). Although CHL is a weak inducer of biofilms, its effect was not impacted by nitrate addition, suggesting that lack of nitrate suppression is not specific to 30S targeting antibiotics. CHL activity is context specific, halting translation when an alanine is in the penultimate position on the nascent polypeptide (Choi et al., 2020; Marks et al., 2016). The macrolides azithromycin (AZI) and erythromycin (ERM) target the 50S subunit at the peptide exit tunnel, and also display context specific inhibition, but for different amino acids than CHL (Kannan et al., 2014). Further, AZI and ERM allow for continued translation of some polypeptides (Kannan, Vázquez-Laslop, & Mankin, 2012), unlike TET and CHL. Despite these similarities, AZI and ERM differ in charge, which affects their uptake and consequent potentiation by bicarbonate (Farha et al., 2020). The biofilm response to sub-MIC AZI and ERM was unaffected by nitrate addition, indicating that nitrate insensitivity is likely not determined by context-specific inhibition, leaky translation, or charge-dependent changes in antibiotic uptake. In our attempts to break the rule of nitrate insensitivity by translation inhibitors, we tested spectinomycin (SPEC), an aminocyclitol antibiotic that targets the 30S subunit and inhibits translocation of tRNAs (Wilson, 2009). Although SPEC has a different mechanism of action from the other inhibitors tested, the biofilm stimulation response to this antibiotic was also insensitive to nitrate. Finally, we tested solithromycin (SOL), belonging to the ketolide subclass of macrolides that – unlike most in the class – is bactericidal rather than bacteriostatic (Svetlov, Cohen, Alsuhebany, Vázquez-Laslop, & Mankin, 2020; Svetlov, Vázquez-Laslop, & Mankin, 2017). Unfortunately, SOL was a markedly worse biofilm inducer than AZI or ERM, so the window for nitrate suppression was narrower, but nonetheless nitrate failed to suppress biofilm stimulation. These data suggest that the effect of nitrate does not bisect with the pathway leading to biofilm stimulation due to inhibition of translation. Given the relatively poor biofilm stimulation by SOL compared to the bacteriostatic macrolides, we tested biofilm stimulation by aminoglycosides, bactericidal translation inhibitors that target the 30S subunit. Strikingly, sub-MIC amikacin, streptomycin, and gentamicin caused little to no biofilm stimulation (**Figure S5**). SPEC, which stimulates biofilm well, is structurally similar to aminoglycosides and binds at the same site on the ribosome, but is bacteriostatic. This result suggests that cidality and translation inhibition may have opposing impacts on the cellular metabolic cues that lead to biofilm stimulation, and a bactericidal translation inhibitor might fail to induce biofilm stimulation because it drives metabolism simultaneously in two incompatible directions.

### Sub-MIC bactericidal antibiotics cause a spike in metabolic activity

To visualize the effects of sub-MIC antibiotics on respiration, we used a disc diffusion assay to generate a gradient of antibiotic concentrations. To monitor changes in cellular respiration we used a tetrazolium dye, MTT (3-(4,5-dimethylthiazol-2-yl)-2,5-diphenyltetrazolium bromide), that is reduced intracellularly by NADH into a crystalline formazan dye (**Figure 6a**) (V. Berridge, Herst M., & Tan S., 2005). Formazan dye deposition (indicated by a dark ring and drop in the pixel intensity) clearly localized in the periphery of the zone of inhibition for CEF and GENT, but not TET (**Figure 6bc**). Thus, the bactericidal antibiotics CEF and GENT increase respiration in the sub-MIC zone, whereas the bacteriostatic antibiotic TET does not.

**Figure 6.**
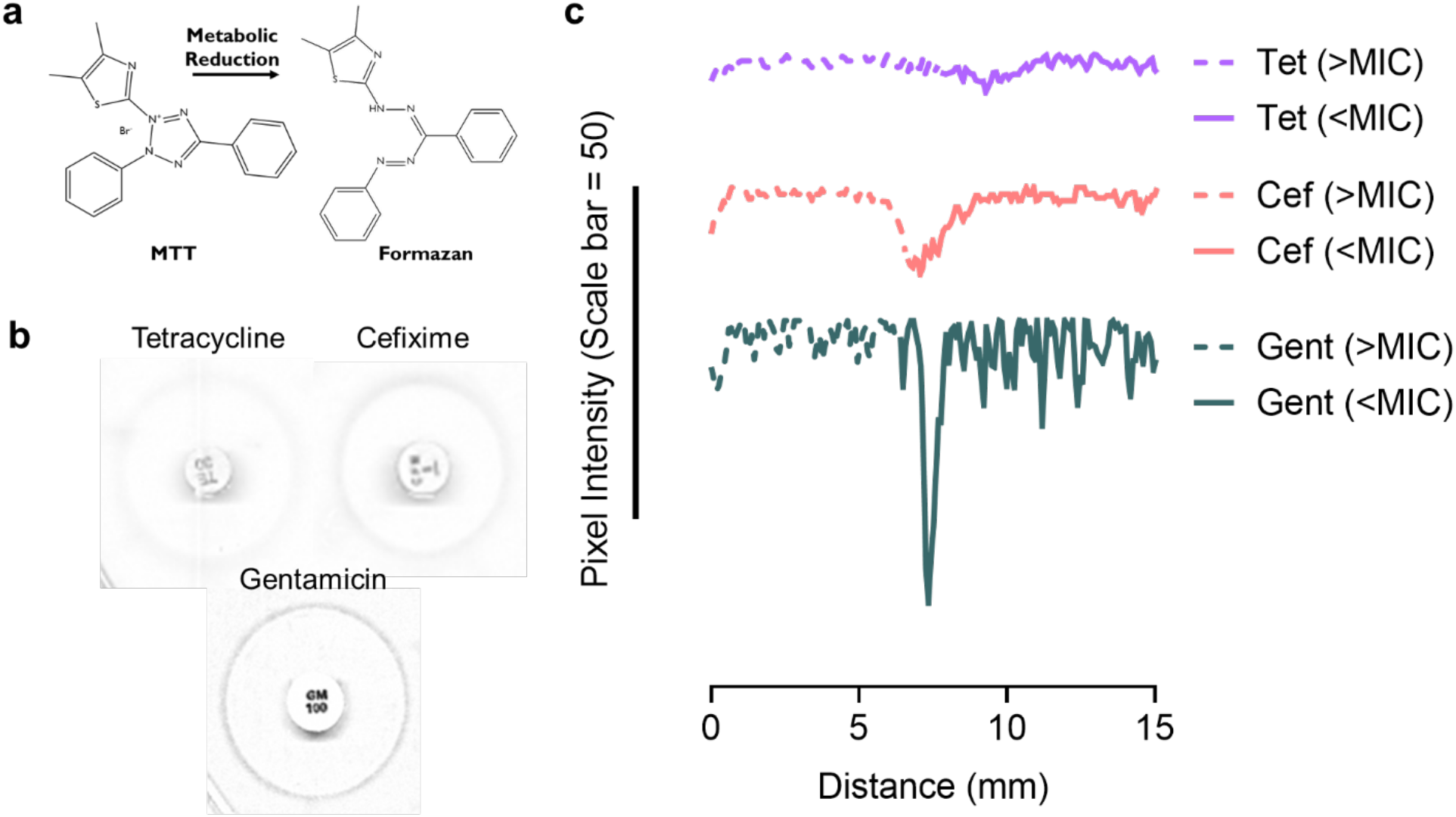
Spatial localization of increased metabolism in the sub-MIC zone. **a)** Reduction of the tetrazolium salt MTT into a formazan dye by cellular metabolism. **b)** Images of the zone of inhibition on the MTT agar plates for TET, CEF, and GENT that were used for quantification. **c)** Pixel intensity as a function of distance (in centimeters) from the edge of an antibiotic disk. The zone of inhibition (>MIC) is marked by a dashed lines and extends from the edge of the disk to the point where growth became visible. The sub-MIC zone (<MIC) is marked by solid lines and extends to the edge of the measured region. Data for TET, CEF, and GENT are indicated by purple, red, and green lines, respectively. To better visualize each line, TET and CEF data points were nudged by 40 and 20 units up the y-axis, respectively, and a scale bar representing 50 units is shown. The graphs are representative of three biological replicates.

## Discussion

### Sub-MIC antibiotics drive biofilm stimulation by changing metabolic activity

Using a high-throughput biofilm stimulation screen of the Keio collection, we identified sub-MIC antibiotic-induced metabolism and respiration perturbations as central consequences leading to the subsequent spike in biofilm levels. A connected network of mutants in the lower TCA cycle and respiration were impaired in the biofilm stimulation response. The Arc and Cpx systems that sense respiratory quinone redox state and envelope stress, respectively, were also required for biofilm stimulation (**Figure 7**). For some antibiotics, the stimulation response was suppressed in a dose- and respiration-dependent manner by the alternate electron acceptor, nitrate. Strikingly, nitrate supplementation had little to no effect on biofilm stimulation by sub-MIC translation inhibitors. Finally, increased metabolic activity colocalized with sub-MIC antibiotic concentrations of bactericidal compounds.

**Figure 7.**
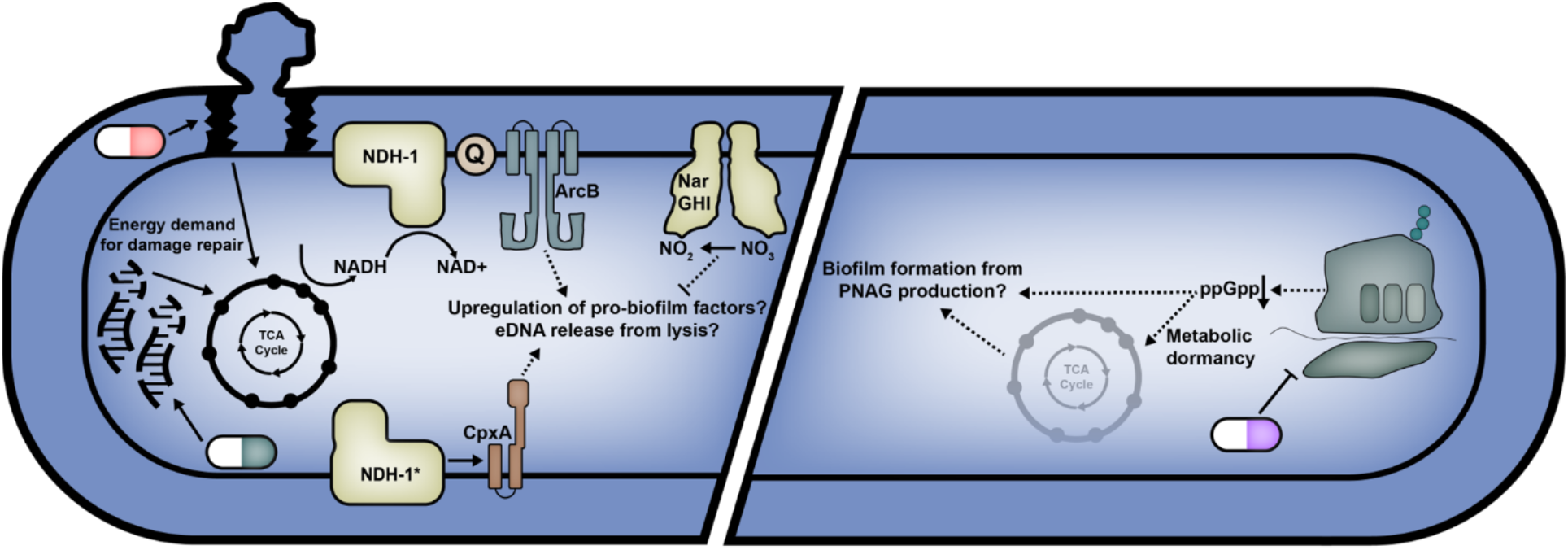
Model of the path from antibiotic stress to biofilm formation. Repairing damage from bactericidal (e.g., cell wall and DNA targeting) antibiotics increases energy demand and consequently flux through the TCA cycle and aerobic respiratory chain. Changes in respiratory chain flux affect the redox status of the quinones (Q) which affects the activity of ArcB (Alvarez et al., 2013). Inner membrane stress from NDH-1 (complex I) overproduction (indicated by *) activates the Cpx system (based on the Guest et al. (2017) model). Cpx and Arc signalling leads to increased biofilm formation through upregulation of biofilm components or increased lysis and eDNA release. Reduction of nitrate by NarGHI suppresses biofilm stimulation. Incorporating our data into the existing model from Boehm et al. (2009), we propose that translation inhibitors induce metabolic dormancy and biofilm formation through reductions in ppGpp.

### Biofilm stimulation provides functional context for antibiotic induced metabolic perturbations

We set out to find the common elements in *E. coli* physiology that are affected by antibiotics and necessary for biofilm stimulation; re-examining the literature through the lens of our data shows that metabolic and respiratory flux were prime candidates for this role. Metabolic and respiratory changes caused by bactericidal antibiotics are well characterized, particularly by the Collins group (Belenky et al., 2015; Dwyer et al., 2014; Lobritz et al., 2022, 2015; Mathieu et al., 2016; Stokes et al., 2019). Their studies suggest that the lethality of bactericidal antibiotics is driven in part by increased flux through oxidative phosphorylation as a non-optimal response to antibiotic damage. Indeed, peptidoglycan synthesis under beta-lactam stress draws more metabolites from glycolysis, causing ROS production (Kawai et al., 2019). It is curious then, given the time scale for co-existence of bacteria and antibiotics, that bacteria lack the means to circumvent the harmful increase in flux, although the selective pressure for this may be too low to drive evolution. Based on our data, we propose instead that this increased metabolic flux may drive an advantageous response by stimulating biofilm formation and providing protection of the population against further antibiotic assault.

Similarly, a network of metabolic genes and the ArcA/B system were implicated in playing an adaptive role by promoting stress-induced mutagenesis (Al Mamun et al., 2012). The Arc and the Cpx systems were also implicated in antibiotic cidality (Kohanski, Dwyer, Wierzbowski, Cottarel, & Collins, 2008). The Collins group found that only bactericidal antibiotics drive increases in metabolic flux, whereas we found that biofilm stimulation occurs in response to both bactericidal and bacteriostatic antibiotics. We showed that the electron acceptor nitrate suppressed biofilm stimulation for the bactericidal cell wall and DNA synthesis-targeting antibiotics CEF and NOVO, but not for the translation inhibitors TET, CHL, AZI, ERM, SPEC, and SOL. This finding suggests that translation inhibitors – unlike other antibiotics – may converge on the biofilm stimulation phenotype through different effects on metabolism, supported by our data showing the lack of MTT dye reduction by cells growing in sub-MIC TET. For example, relative to untreated cells, CHL treatment reduces oxygen consumption rate (Dwyer et al., 2014), suggesting that bacteriostatic antibiotics reduce, rather than increase, demand for electron acceptors. The dampening of metabolic demand by bacteriostatic antibiotics is in line with previous work on *E. coli* biofilm stimulation by the bacteriostatic translation inhibitors TET and CHL, where biofilm stimulation by these drugs involves derepression of PNAG biosynthesis through suppression of ppGpp production (Boehm et al., 2009). Notably, a ppGpp null mutant has decreased expression of many central metabolism genes (Traxler et al., 2008). From this, we speculate that bacteriostatic and bactericidal antibiotics may converge on biofilm formation through opposite demands on the central metabolism and respiration cues that signal biofilm stimulation. This hypothesis would explain why bactericidal translation inhibitors were poor biofilm stimulators. The uncoupling of growth inhibition and biofilm stimulation by nitrate could prove to be a useful phenotype for future studies on the direct and downstream effects of bactericidal antibiotics.

### Toward the origin and evolution of biofilm stimulation

Many chemically- and mechanistically-distinct antibiotics can induce biofilm formation in a range of bacterial species (Boehm et al., 2009; Hoffman et al., 2005; Jin et al., 2020; Kaplan et al., 2012; López, Fischbach, Chu, Losick, & Kolter, 2009; Nguyen et al., 2014; M. R. Ranieri et al., 2018; Tahrioui et al., 2019; Townsley & Shank, 2017). Certain antibiotics inhibit biofilm formation, but these are usually strain or drug-specific effects rather than a generalized response (Ichimiya et al., 1996; Kimura et al., 2017; Moon, Weber, & Feldmana, 2017). The variety of antibiotics and species demonstrating similar biofilm phenotypes suggest a common, underlying mechanism of response to antibiotic stress, rather than reaction to the direct detection of toxic compounds. Boehm *et al*. showed this elegantly with an *E. coli* mutant that required the aminoglycoside streptomycin to grow, where biofilm production increased in response to decreasing concentrations of streptomycin (Boehm et al., 2009). Previous work on biofilm stimulation focused on the roles of specific signalling molecules like cyclic-di-GMP (Boehm et al., 2009; Hoffman et al., 2005), ppGpp (Boehm et al., 2009), and quorum sensing molecules (López et al., 2009; Tahrioui et al., 2019), or eDNA release mechanisms (Jin et al., 2020; Kaplan et al., 2012; Mlynek et al., 2016; Tahrioui et al., 2019; Yu, Hallinen, & Wood, 2018). Release of eDNA and bacterial cell death are critical steps in biofilm formation and have many parallels to programmed cell death in eukaryotes (Bayles, 2007; Turnbull et al., 2016). Metabolic overflow pathways are important for bacterial programmed cell death and subsequent biofilm development in *S. aureus*, in line with our model (Thomas et al., 2014). It is worth further investigating the parallels between antibiotic-induced biofilm formation and programmed cell death in eukaryotes, as this may shed light on common origins of these pathways (Dwyer, Camacho, Kohanski, Callura, & Collins, 2012). For example, the *P. aeruginosa* PQS quorum sensing derivative HQNO causes cell lysis by disrupting cytochrome *bc1* in the respiratory chain (Hazan et al., 2016), while PQS contributes to biofilm stimulation through an unknown mechanism (Tahrioui et al., 2019).

### Biofilm stimulation deepens the connection between respiration status and biofilms

It is clear that metabolic changes are required for antibiotic-induced biofilm formation, but a role for cellular redox state and respiration is well documented in normal biofilm formation as well (Okegbe, Price-Whelan, & Dietrich, 2014; Sporer, Kahl, Price-Whelan, & Dietrich, 2017). Limiting oxidative respiration leads to multicellularity in *Bacillus subtilis* and *S. aureus*, which can be rescued by supplying an alternative electron acceptor (Kolodkin-Gal et al., 2013; Mashruwala, van de Guchte, & Boyd, 2017). These responses are mediated by the sensor kinases KinB and SrrB in *B. subtilis* and *S. aureus*, respectively. In *B. subtilis*, KinB increases biofilm formation by phosphorylating the master sporulation regulator Spo0A, while respiration control of biofilm formation in *S. aureus* depends on sensing of the menaquinone pool oxidation state by the SrrA/B two-component system and increased transcription of *atlA*, encoding an autolysin that releases eDNA. A *S. aureus atlA* mutant is notably unable to respond with increased biofilm formation when exposed to sub-MIC methicillin (Kaplan et al., 2012). In *P. aeruginosa*, colony morphogenesis and biofilm formation are influenced by redox state and extracellular electron exchange from phenazines and a cbb3-type oxidase (Dietrich et al., 2013; Jo, Cortez, Cornell, Price-Whelan, & Dietrich, 2017). We show here that ArcA/B might function similarly to SrrA/B in sensing changes in respiration status and coordinating biofilm formation.

Alternatively, ArcA in *Vibrio cholerae* can be activated by ROS in oxygen-rich conditions that normally inactivate it (Zhou et al., 2021). We have yet to identify the pro-biofilm effector(s) controlled by ArcA/B, and although the focus of this work was on global physiological changes leading to biofilm stimulation, we showed that curli production is dispensable for increased biofilm induced by sub-MIC antibiotics. One possibility for signalling downstream of ArcB is the Cpx system. Arc and Cpx undergo crosstalk under gentamicin stress (Ćudić et al., 2017), however, if crosstalk was responsible for biofilm stimulation, then the presence of ArcB or ArcA alone should be sufficient (either as a phosphodonor or receiver with Cpx), but we did not find this to be true. The Cpx system is involved in downregulation of cellulose production through OmpA, which causes increased or decreased biofilm formation on hydrophobic or hydrophilic surfaces, respectively (Ma & Wood, 2009). The pathway leading from the Arc/Cpx systems to antibiotic-induced biofilm formation is worth investigating in future work.

### Concluding remarks

Aspects of the biofilm stimulation response have been characterized in select organisms with select antibiotics; however, to our knowledge this is the first systems-level approach focusing on essential components of the response across multiple antibiotics. We found that global changes in central metabolism and respiration connect treatment with sub-MIC antibiotics to increased biofilm formation. This work also provides context for recent studies on antibiotic-induced metabolic changes, as such changes may constitute a general strategy for inducing biofilm formation and providing subsequent population-level protection against higher concentrations of antibiotics. If increased metabolism/respiration causes a biofilm-promoting protective response, this work would caution against inducing futile cycles as a potential therapeutic strategy (Adolfsen & Brynildsen, 2015). The biofilm stimulation response has also proved useful in our lab as a hypersensitive screen for otherwise overlooked subinhibitory antibiotic activity (Coles et al., 2022; Nguyen et al., 2012; M. R. M. Ranieri et al., 2019).

The work detailed here suggests new possibilities for adjuvant development, detailing targets that, when inhibited, should suppress biofilm stimulation. Mutations in some of the core metabolic genes identified from our screens (*icd, sucA, nuoM*) that confer antibiotic resistance have been identified in clinical isolates (Lopatkin et al., 2021), so inhibiting these targets might sensitize cells in addition to suppressing biofilm stimulation. Targeting these metabolic enzymes must avoid inhibition of their eukaryotic homologues, but careful design of narrow spectrum inhibitors of bacterial central metabolism is possible (Keohane et al., 2018; Shapiro, Kaplan, & Wuest, 2019). We also showed that nitrate suppresses biofilm stimulation by bactericidal antibiotics. *E. coli* is a facultative anaerobe, and regularly encounters oxygen-limited environments when establishing an infection. Although our experiments were performed with a laboratory strain, if the nitrate suppression phenotype is consistent for pathogenic strains, then antibiotic treatment could be coupled with dietary intake of nitrates to prevent the stimulation of biofilm formation. In a similar sense, changes in pH affect the sensitivity of *E. coli* to certain beta-lactams (Mueller, Egan, Breukink, Vollmer, & Levin, 2019), and bicarbonate re-sensitizes pathogens to AZI (Farha et al., 2020), so manipulating host environmental conditions could be a simple and effective strategy to improve antibiotic efficacy. By characterizing the systems level effects of sub-MIC antibiotics on biofilm formation, we’ve expanded the repertoire of environmental factors and small molecule targets that modulate the response of *E. coli* to antibiotic assault.

## Materials and methods

### Bacterial strains, plasmids, and culture conditions

*E. coli* studies used the parent strain of the Keio Collection, *E. coli* K-12 BW25113 [F−*Δ(araD-araB)567 lacZ4787Δ::rrnB-3 LAM− rph-1 Δ(rhaD-rhaB)568 hsdR514*] (denoted herein as *E. coli*) (Baba et al., 2006). Key mutants selected from the Keio collection were verified with PCR. For overnight cultures, −80ºC stocks were used to inoculate 3 mL of lysogeny broth (LB; Lennox) and incubated at 37ºC for 16 h while shaking at 200 rpm. Subcultures were made in 50:50 (50% LB and 50% phosphate buffered saline) and nitrate (KNO_3_ or NaNO_3_) was supplemented where indicated. Nitrate broth (5g/L peptone, 3g/L meat extract, and 50 mM potassium nitrate) was used for the nitrate reduction assay. Plates for solid media assays were made using 1.5% agar and 50:50 media.

### Peg lid biofilm assay

The peg lid biofilm assays were performed as described previously by Ceri et al., with minor modifications (Ceri et al., 1999; M. R. M. Ranieri et al., 2019). Briefly, overnight cultures were diluted 1:100 into a 3 ml subculture of 50:50, grown for ∼2 hours and normalized to an OD600 of 0.1 (DeNovix DS-11+ Spectrophotometer), then diluted 1:500 in 50:50. Each well of a flat bottom 96-well plate (Nunc) contained 148 μl of inoculated media and 2 μl of antibiotic/vehicle control. As a sterility control, 148 μl of sterile media and 2 μl of vehicle was pipetted into each well of row H. A peg lid (Nunc) was then set in the plate which was sealed with parafilm. Plates were incubated in a humidity-controlled chamber at 37ºC for 18 h while shaking at 200 rpm. After, the peg lid was removed and planktonic growth was measured (OD600, Multiskan Go – Thermo Fisher Scientific). The lid was then washed in phosphate-buffered saline (PBS: 137 mM NaCl, 2.7 mM KCl, 10 mM Na_2_HPO_4_, 1.8 mM KH_2_PO_4_, pH 7.4) and stained in 0.1% (wt/vol) crystal violet for 15 min. Excess crystal violet was removed by 3 separate washes in Milli-Q water and the lids were left to dry. The crystal violet attached to each peg was solubilized in 200 μl of 33% acetic acid (vol/vol) and absorbance was measured at 600 nm (A600, Multiskan Go – Thermo Fisher Scientific). Raw values were plotted as a percent of the vehicle control and statistical analysis was performed using Prism (GraphPadv9.3.1). Each condition was tested in technical triplicate.

### High throughput Congo Red binding screen

For the Keio Collection screen, we poured 25 mL of media (50:50 broth, 1.5% agar, 40 μg/ml Congo Red) supplemented with one of 125 μg/ml novobiocin, 0.6 μM cefixime, or 1 μg/ml tetracycline into rectangular plates. From a source plate, colonies were pinned on the antibiotic-containing plates and once on a vehicle control plate at 1536 colony density using the ROTOR HDA (Singer Instruments). This experiment was done in duplicate using two different source plates. Plates were incubated for 24 h at 37 ºC, then scanned (Epson Perfection V750) and analyzed with a custom FIJI (ImageJ) program detailed in Appendix I (French et al., 2016). Data from colonies was normalized by plate position by taking the interquartile mean of each column or row, then dividing each value in that column or row by the interquartile mean. The normalized Congo Red values were divided by the normalized growth values to generate the enrichment value. Mutants with normalized growth below 0.2 were excluded from the hit list.

### Measuring nitrate reduction

Nitrate reduction was measured as previously described with some modifications to accommodate a 96-well format (Buxton, 2011; Griess, 1858). Briefly, 96-well plates were set up in the same manner as the peg-lid biofilm assay with serial dilutions of antibiotics across the rows, except peg-lids were not added prior to incubation. Strains were sub-cultured in nitrate broth prior to adding to the plate, then transferred in 148 μL of nitrate broth +50mM KNO_3_ for each well then grown for 18 hours at 37 ºC while shaking, and OD600 was read with a spectrophotometer. Five μL each of nitrate reagent A and B (Sigma) were added to a new 96-well plate, and 100 μL of the culture was added. Absorbance was read at 546 nm immediately after to minimize dye degradation. Raw values were plotted as a percent of the vehicle control and statistical analysis was performed using Prism (GraphPadv9.3.1). Each condition was tested in technical triplicate.

### Disk diffusion MTT reduction

Plates made of 50:50 LB and 1.5% agar were supplemented with 0.04% MTT and dried in a biosafety cabinet. Next, a 10 mL soft LB agar overlay was poured over the plates with the same concentration of MTT, 0.6% agar, and 100 μL of 0.3 OD_600_ *E. coli* culture and dried. Once dry, antibiotic disks (Oxoid) were placed on the soft agar overlay with sterile forceps and the plates were incubated for 18 h at 37 ºC. The plates were then imaged with a scanner (Canon). ImageJ was used to analyze the resulting images. Prior to analysis, the MTT plate images were split into their RGB colour channels, the green channel was isolated, and the background was subtracted followed by running the ‘enhance contrast’ function. The ‘plot profile’ function was used to quantify pixel intensity from the edge of the antibiotic disk to the sub-MIC zone just beyond the boundary of the zone of inhibition. Pixel intensity was plotted as a function of distance from the edge of the disk. The data from both plates were graphed using Prism (GraphPadv9.3.1) to compare the pixel intensity profiles of each antibiotic.

## Supplemental Tables

**Table S1.**
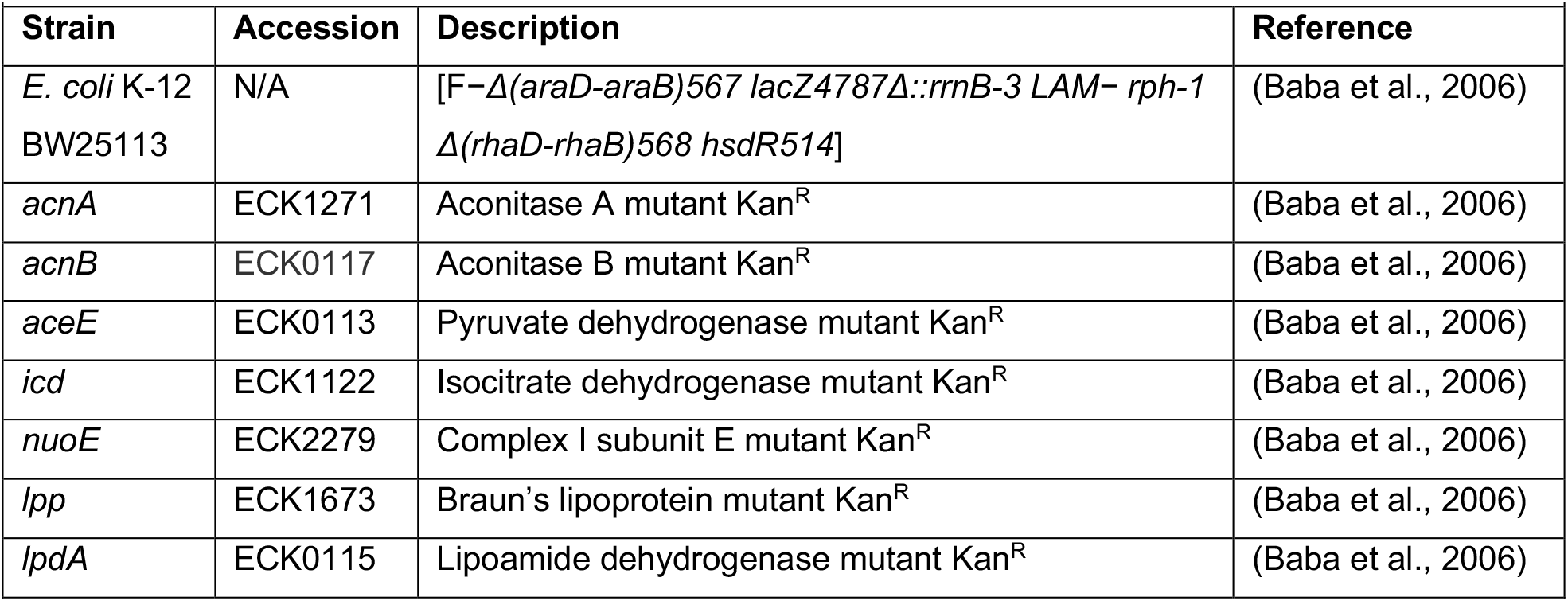
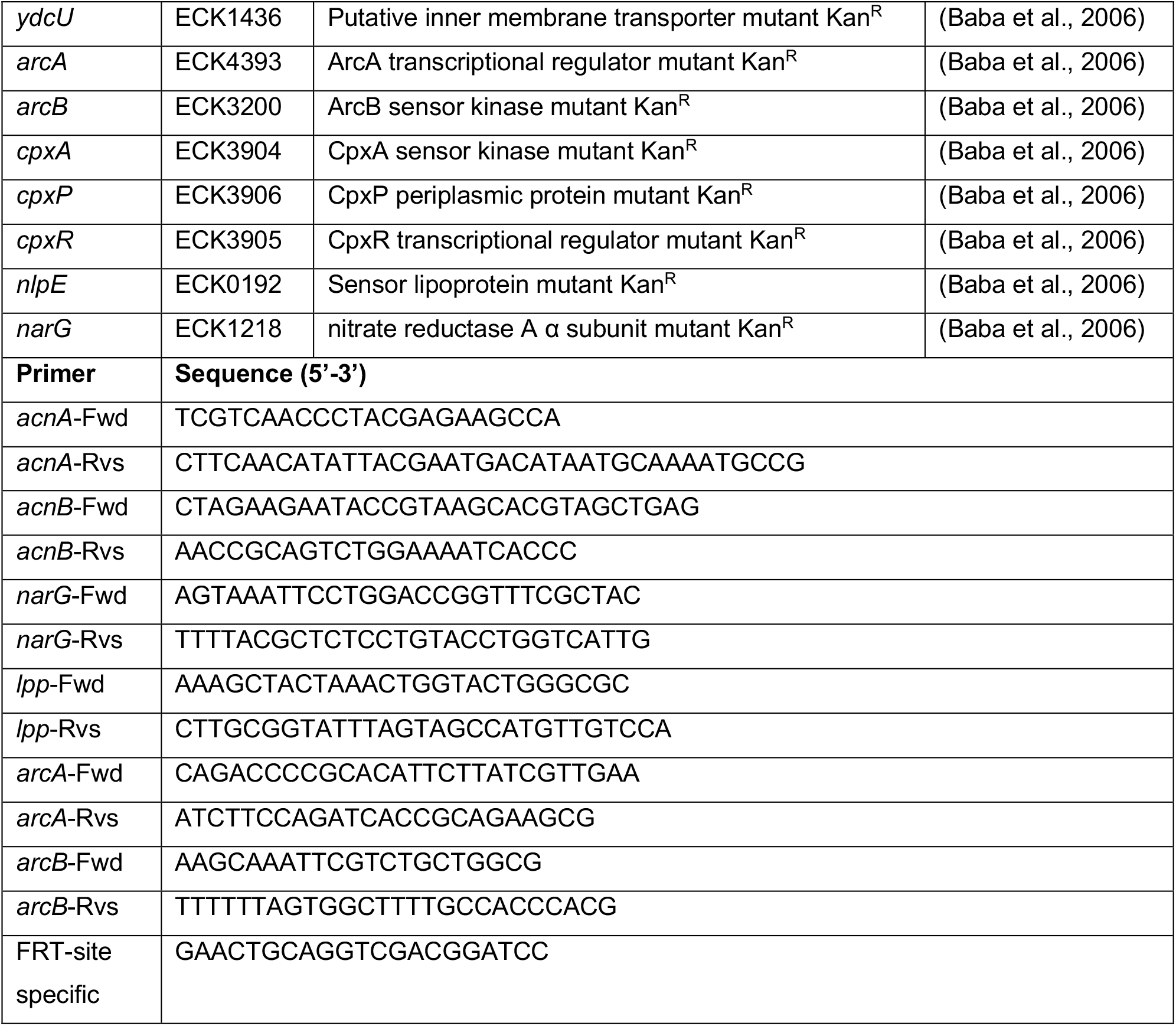
Strains and Primers used in this study.

**Table S2.**
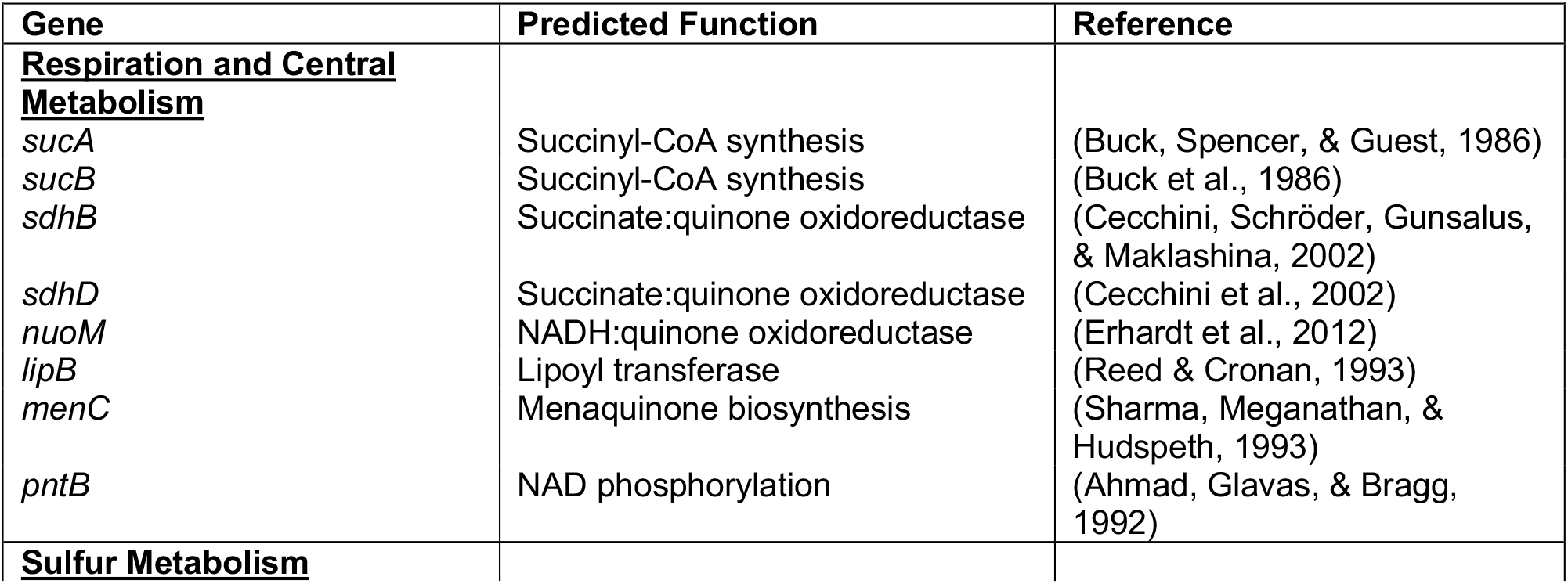
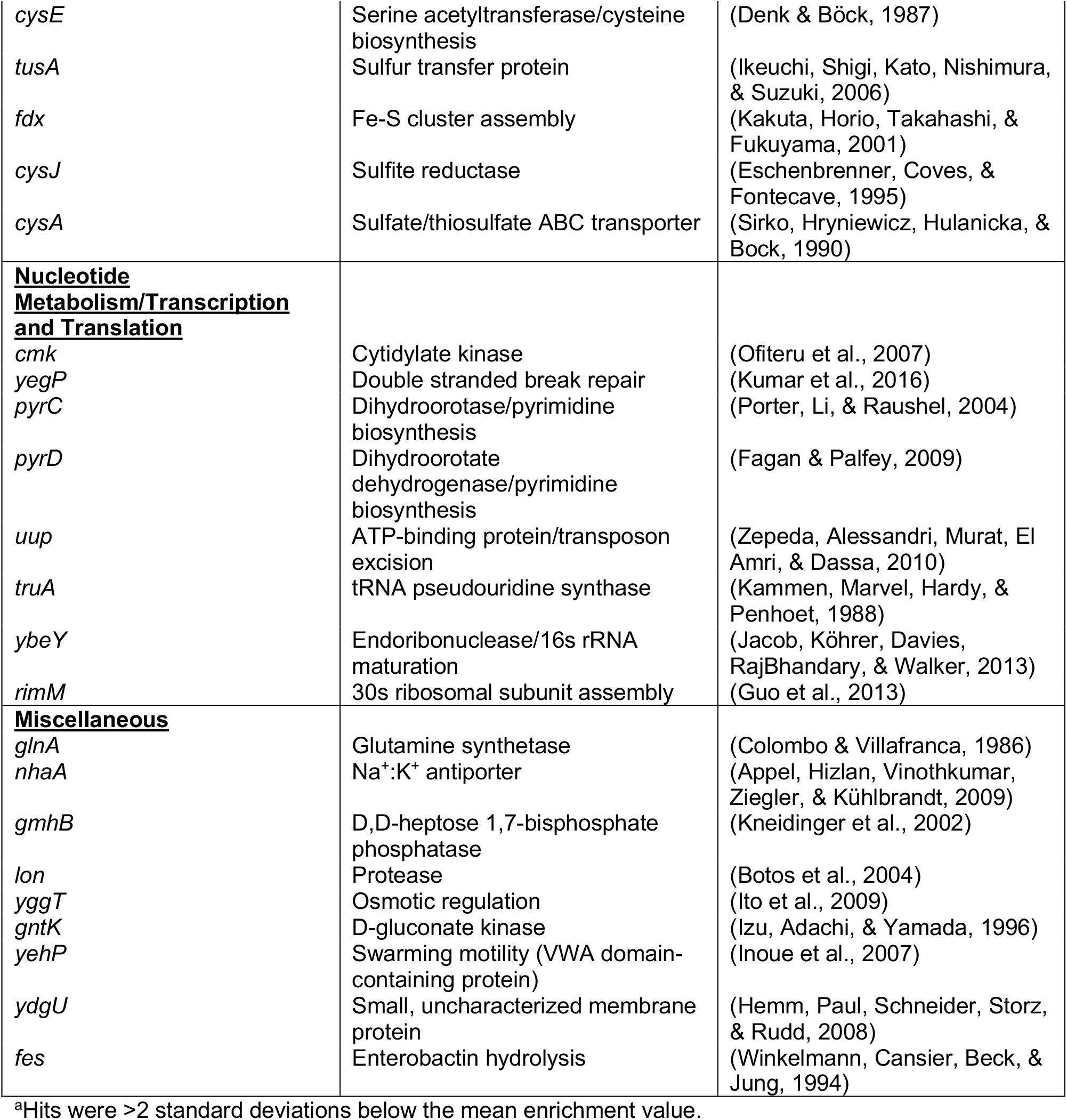
Keio screen hits^a^ and predicted function common to three sub-MIC antibiotics.

## Supplemental Figures

**Supplementary Figure S1.**
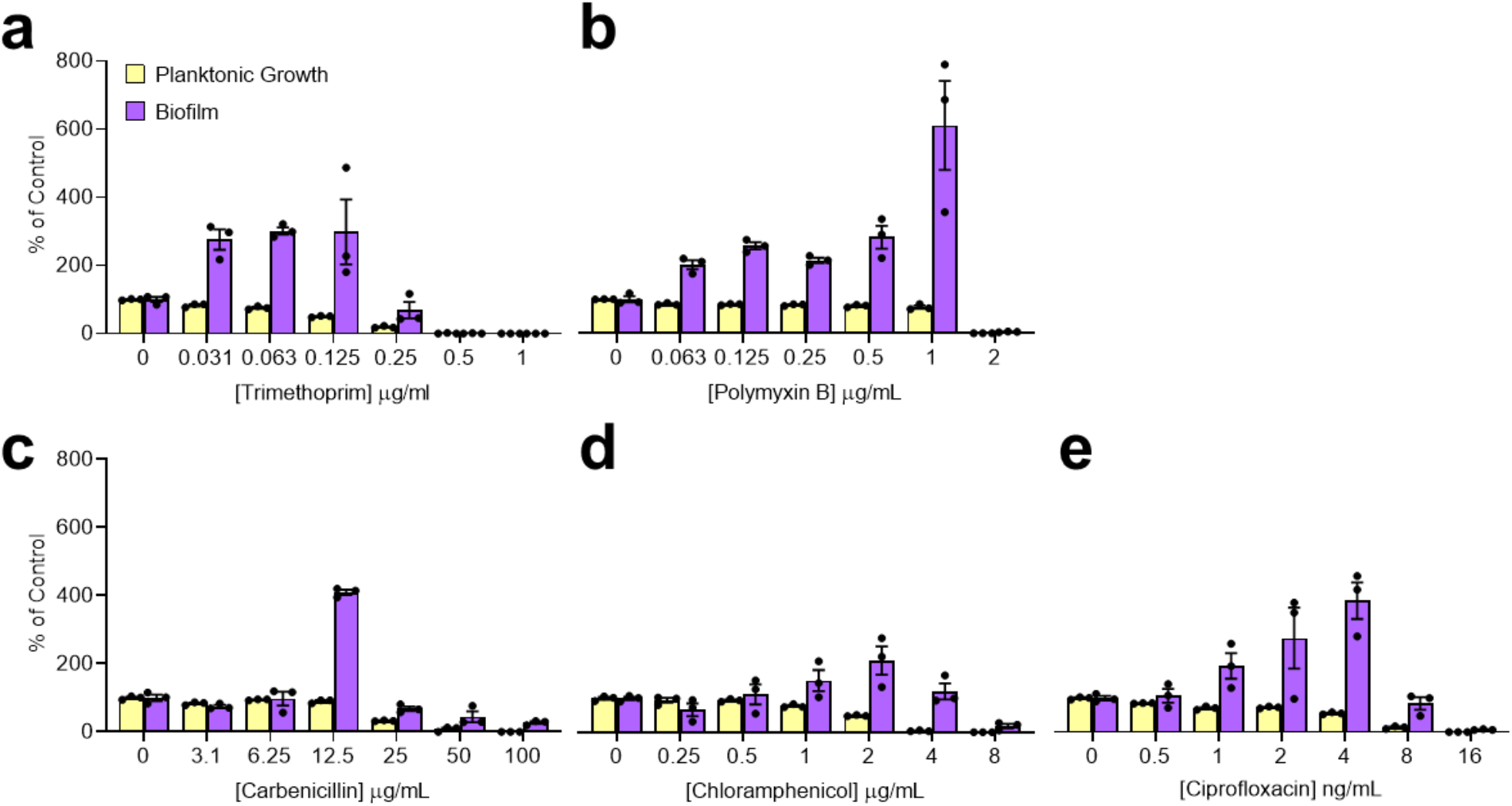
Multiple antibiotics stimulate *E. coli* biofilm formation. The effect of increasing concentrations of **a)** trimethoprim, **b)** polymyxin B, **c)** carbenicillin, **d)** chloramphenicol, and **e)** ciprofloxacin on *E. coli* K12 biofilm stimulation was measured using a peg lid assay. All values are relative to the untreated vehicle control for each respective antibiotic. Circles show the value of each technical replicate of the triplicate, columns show the mean of each replicate, and the bars show the standard error of the mean. Data points are shown as a percent of control, and the y-axis scale on the left applies to all the graphs.

**Supplementary Figure S2.**
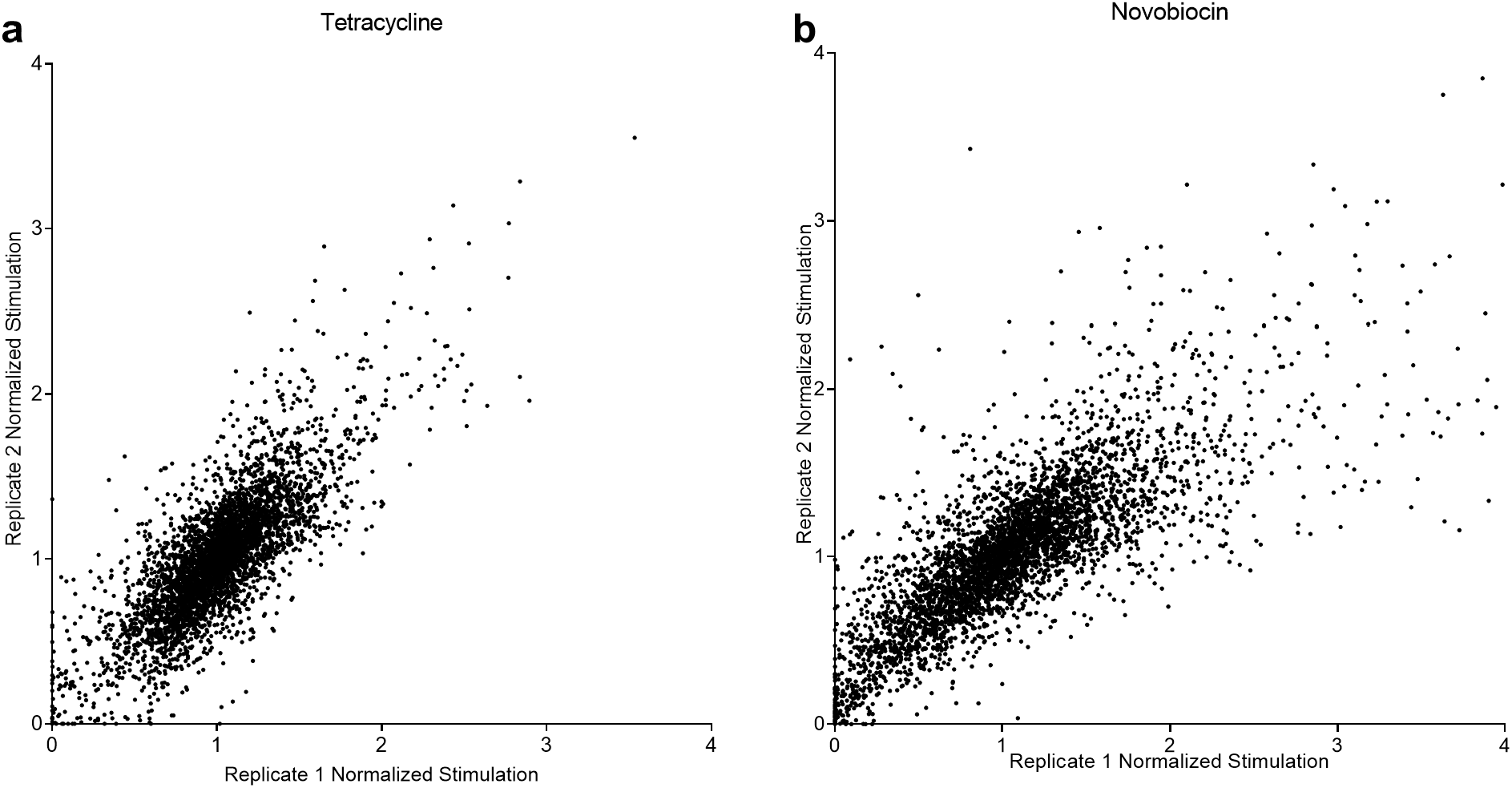
Biofilm stimulation response to TET and NOVO for the Keio collection. Screening results for TET and NOVO shown in a replica plot with extremely high stimulation data points (above 4) removed to better visualize low stimulation data. Screening was done in technical duplicate and data from each replicate were plotted on the x and y-axis.

**Supplementary Figure S3.**
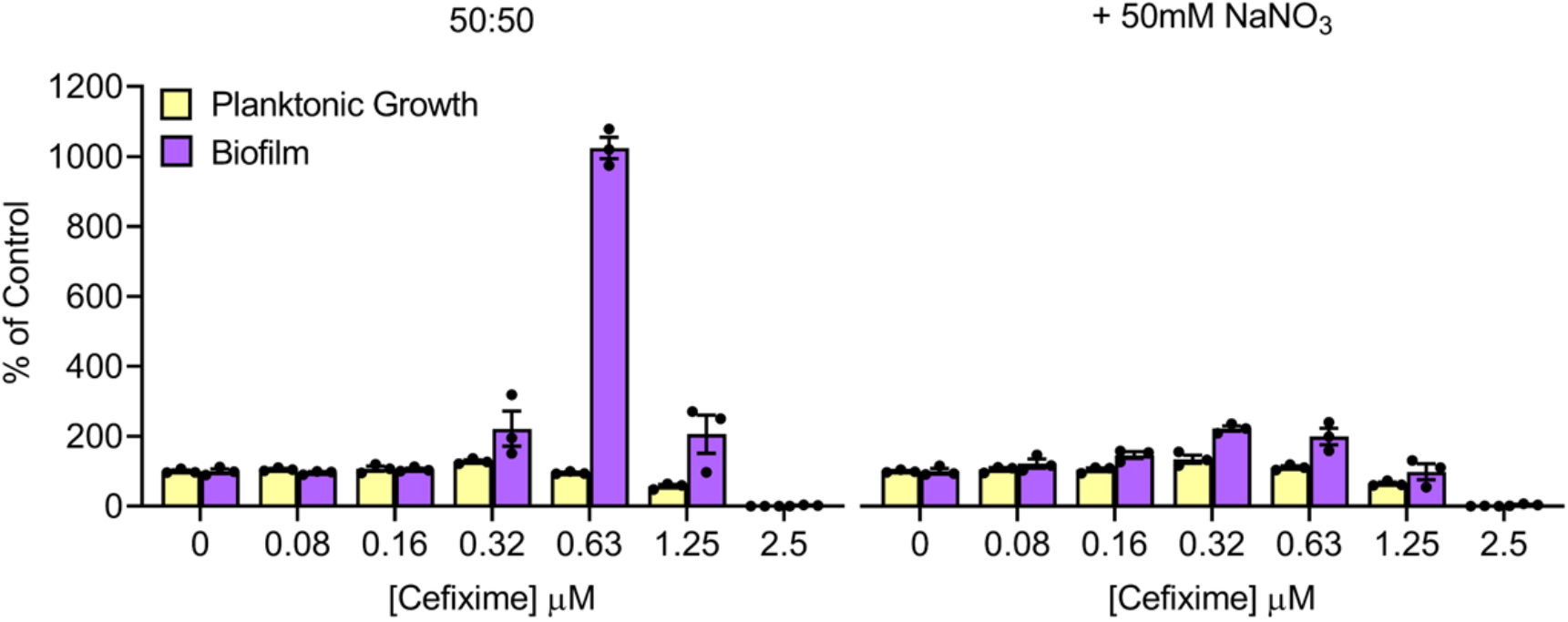
Sodium nitrate supplementation suppresses biofilm stimulation. The effect of adding 50 mM sodium nitrate on biofilm stimulation was determined by measuring biofilms with a peg lid assay across a range of CEF concentrations. Data points show the growth and biofilm of each well relative to the respective untreated vehicle control (as a percent of control). The columns show the mean of each technical triplicate, and the error bars show the standard error of the mean. The y-axis on the left applies to both graphs. The graphs are representative of 3 biological replicates.

**Supplementary Figure S4.**
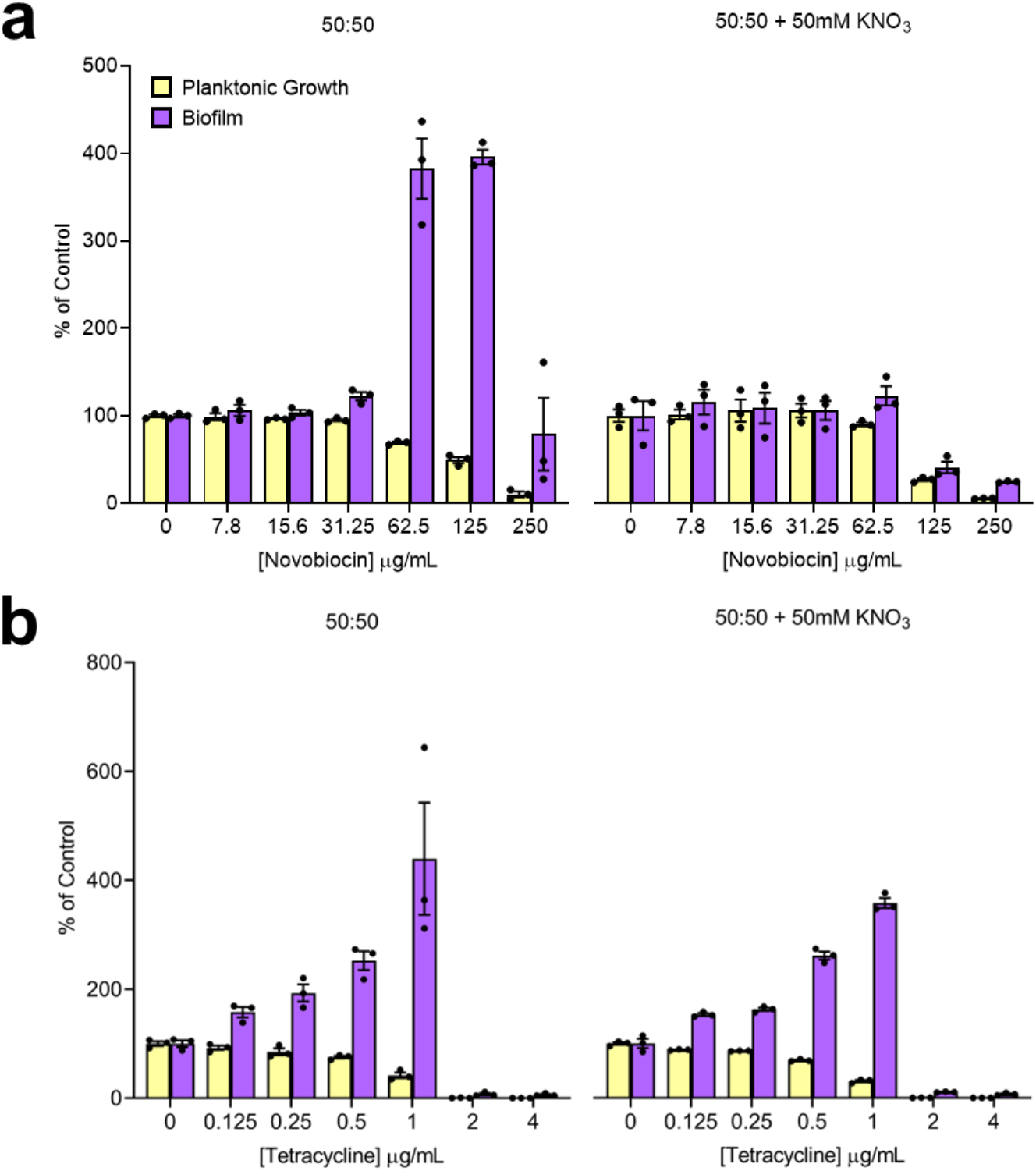
Effects of providing nitrate on biofilm stimulation by novobiocin and tetracycline. Peg lid biofilm assays were done with increasing concentrations of **a)** NOVO or **b)** TET and either no nitrate (left) or 50 mM nitrate (right). Growth and biofilm formation are shown as a percent of the vehicle control. Error bars represent the standard error of the mean. Circles show the value of each technical replicate of the triplicate, columns show the mean of each replicate, and the bars show the standard error of the mean. The graphs are representative of 3 biological replicates.

**Supplementary Figure S5.**
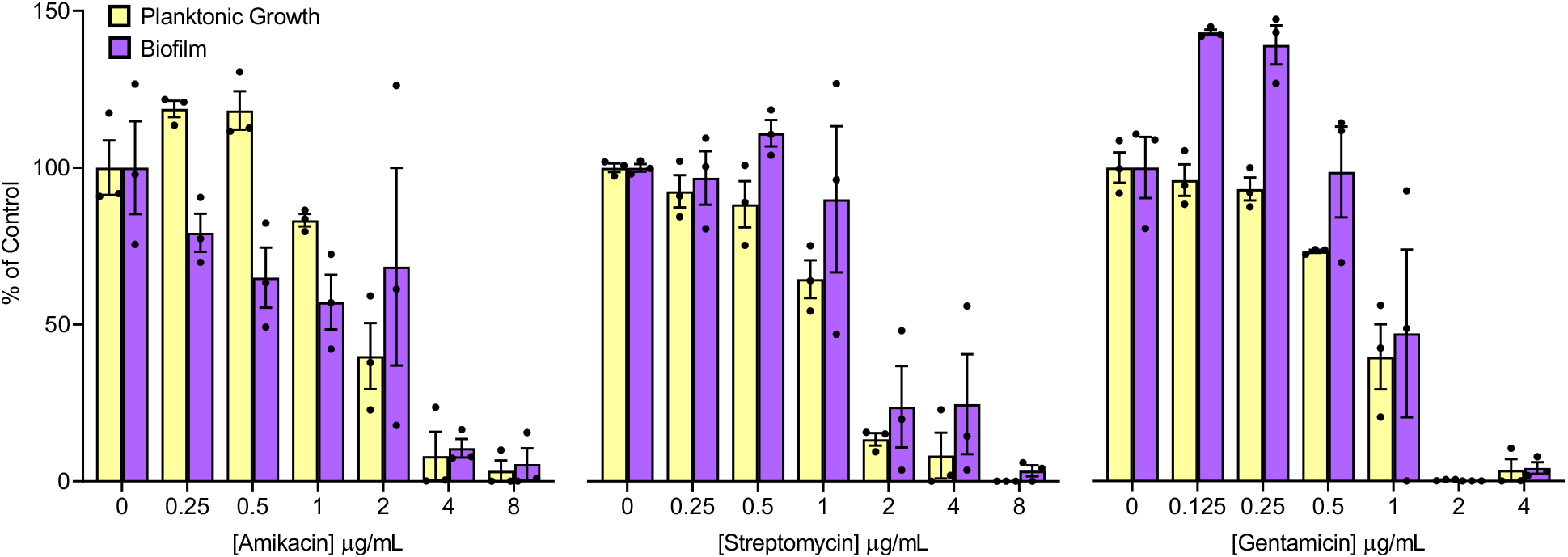
Aminoglycosides are poor biofilm-stimulating antibiotics. Peg lid biofilm assays were done with increasing concentrations of amikacin, gentamicin, or streptomycin. Growth and biofilm formation are shown as a percent of the vehicle control. Circles show the value of each technical replicate of the triplicate, columns show the mean of each replicate, and the bars show the standard error of the mean. The graphs are representative of 3 biological replicates.

## Appendix I

Annotated ImageJ macro for automated colony analysis (.ijm file)

Excel file containing raw and normalized screening data (.xlsx file)

## Competing Interests

The authors have no competing interests to declare.

## Notes

### Competing Interest Statement

The authors have declared no competing interest.

